# A mathematical model of whole-body potassium regulation: Global parameter sensitivity analysis^*^

**DOI:** 10.1101/2023.11.10.566654

**Authors:** Melissa M. Stadt, Anita T. Layton

## Abstract

Potassium (K^+^) is an essential electrolyte that is tightly regulated by various complex physiological mechanisms. In this study, we analyze a mathematical model of whole-body K^+^ regulation to investigate the sensitivity of different model outcomes to parameter values. We used the Morris method, a global sensitivity analysis technique, to evaluate the impact of the parameters on both steady state results and transient simulations during a single-meal. Our results shows that the most influential parameters and processes depend on what you are measuring. Specifically, steady state results relied primarily on parameters that were involved in kidney function, while transient results relied on hormonal feedback mechanisms. This study shows that our mathematical model of whole-body potassium regulation captures known physiological function of potassium regulation despite a large number of uncertain parameters.

**MSC codes:** 68Q25, 68R10, 68U05

## 1. Introduction

Potassium (K^+^) is necessary for normal cellular function, including the resting cellular-membrane potential and the propagation of action potentials in neuronal, muscular, and cardiac tissue. K^+^ also plays an vital role in hormone secretion and action, vascular tone, systemic blood-pressure control, gastrointestinal motility, acid–base homeostasis, glucose and insulin metabolism, mineralocorticoid action, renal concentrating ability, and fluid and electrolyte balance [7]. As such, plasma K^+^ concentration ([K^+^]) is strictly regulated and maintained within a narrow range of 3.5 to 5.0 mmol/L, by multiple mechanisms that together attain K^+^ homeostasis [12].

Total body K^+^ content is maintained by the kidneys, which match K^+^ intake with K^+^ excretion [4, 8]. Because adjustments in renal K^+^ excretion happen on a time-scale of several hours, changes in extracellular [K^+^] are initially buffered by movement of K^+^ into or out of skeletal muscle cells. That regulation of the distribution of K^+^ between the intracellular and extracellular compartments, referred to as internal K^+^ balance, occurs during a shift in plasma [K^+^], e.g., immediately after a K^+^-rich meal. Some of the interactions among the multitude of physiological systems and processes involved in K^+^ regulation remain to be incompletely understood. For example, which processes are more critical in maintaining K^+^ at steady state versus immediately after a meal?

To answer this question, we analyze a computational model of whole-body K^+^ regulation developed in Ref. [19]. The model represents the intra- and extracellular fluid compartments as well as a detailed kidney compartment. The model simulates (i) the gastrointestinal feed- forward control mechanism, (ii) the effect of insulin and (iii) aldosterone (ALD) on Na^+^-K^+^- ATPase K^+^ uptake, and (iv) ALD stimulation of renal K^+^ secretion. We conduct parameter sensitivity analysis on the model to investigate the impact of different regulatory mechanisms on K^+^ homeostasis in steady state and transient states.

Like many models of complex physiological systems, the present K^+^ model involves a large number of parameters. Sensitivity analysis can be used to elucidate the relationship between parameters and mechanisms to the results of the model [16, 17, 18]. This method has been applied to many applications of biological systems including blood clotting, cancer modeling, biochemical models, epidemiology, and more [2, 6, 9, 10, 17, 21]. There are two types of sensitivity analysis: local and global methods. Local methods examine the sensitivity of the model inputs at one specific point in the input space by changing parameters one at a time and evaluating the change in the model output. While this is a popular technique, because it is relatively easy to implement and compute, it has a number of disadvantages, such as neglecting how parameters interact with other parameters, and being heavily dependent on the initial parameter set. Global methods take sensitivities at multiple points and are considered better for nonlinear systems. However, global methods are much more computationally expensive. For more details about local and global sensitivity analysis, see Ref. [16]. In this study we use the Morris method to evaluate the global sensitivity of the K^+^ homeostasis model.

## 2. Methods

We conduct a sensitivity analysis by applying the Morris method (see Section 2.2) to the mathematical model of whole-body K^+^ regulation described in Section 2.1. We analyze the impacts of parameter values on (i) the steady state solution and (ii) single-meal simulations with varying K^+^ amounts (Section 2.3).

### 2.1. Mathematical model of whole-body potassium regulation

We apply the Morris method to our published mathematical model (Ref. [19]). The model is a compartmental ordinary differential equation (ODE) model, which incorporates feedforward and feedback effects representing the regulation of K^+^ levels in a man. A model schematic is shown in Fig. 1. The following sections provide a brief description of the model equations. For a more comprehensive description of the model’s development, see Ref. [19].

**Figure 1.**
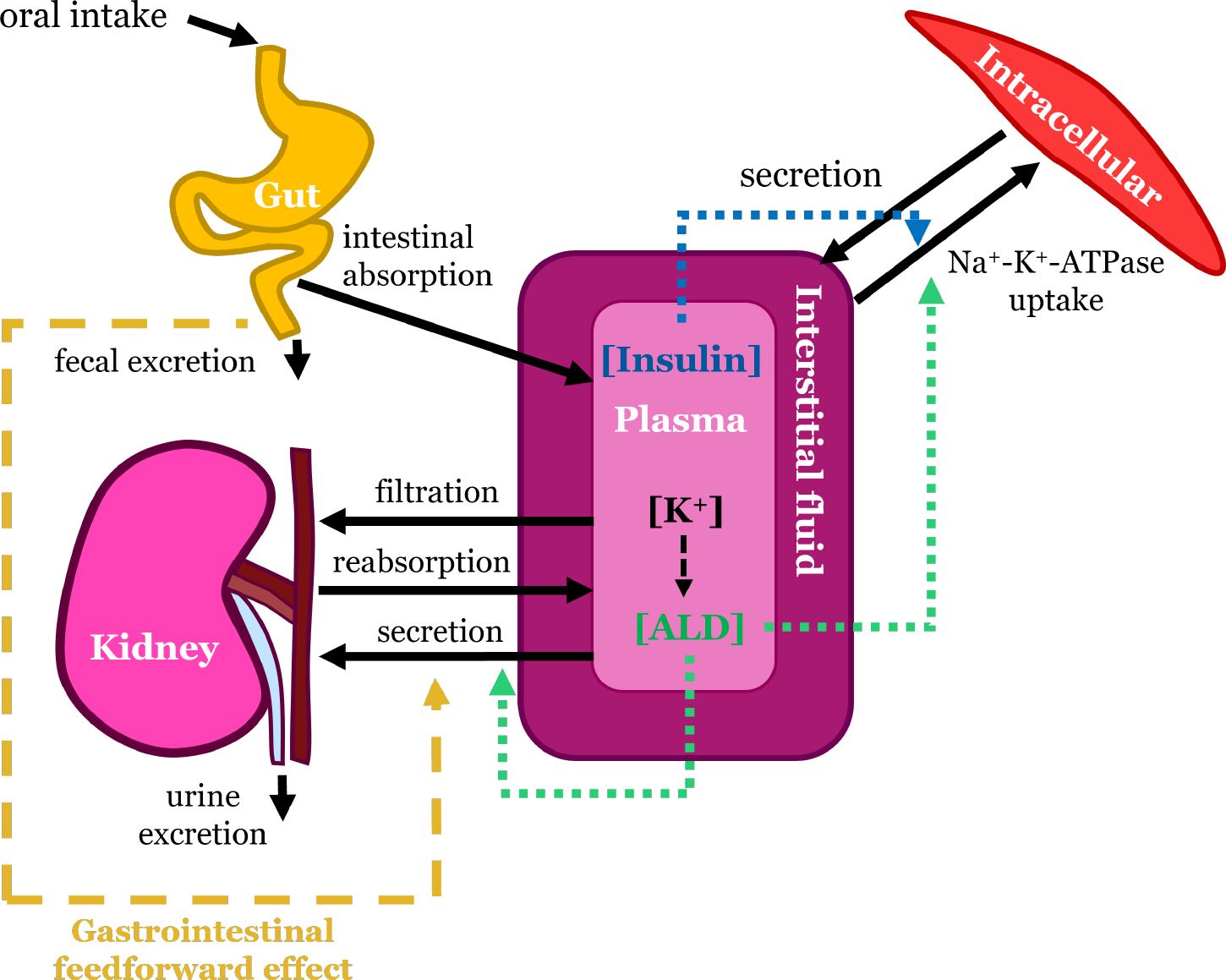
Baseline schematic of K^+^ regulation. Black arrows represent transport of K^+^ between the compartments. Green, light blue, and yellow arrows represent the stimulating effects of aldosterone, insulin, and the gastrointestinal feedforward mechanism, respectively. Note that K^+^ is assumed to diffuse between the interstitial fluid and plasma which make up the extracellular fluid. ALD: aldosterone

#### 2.1.1. Internal K^+^ balance

Internal K^+^ is divided into 4 compartments: gastrointestinal and hepatoportal circulation (denoted by *M*_Kgut_), plasma [K^+^] (denoted by *K*_plasma_), interstitial fluid [K^+^] (*K*_inter_), and intracellular [K^+^] (*K*_IC_).

We determine *M*_Kgut_ by oral K^+^ intake minus K^+^ leaving the hepatoportal circulation:

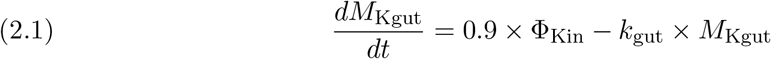

where Φ_Kin_ is K^+^ intake and *k*_gut_ is a parameter that determines the rate of K^+^ delivery to the plasma. Similarly *K*_plasma_ is given by the amount of K^+^ coming into the plasma minus the amount leaving the plasma:

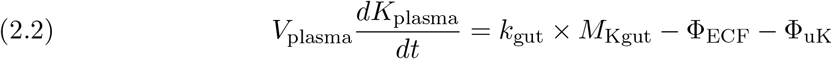

where *V*_plasma_ is plasma volume, Φ_uK_ denotes urine K^+^ excretion (described in Section 2.1.2); Φ_ECF_ denotes the diffusion of K^+^ from the blood plasma to the interstitial space and is given by

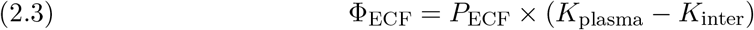

where *P*_ECF_ is a permeability parameter. Interstitial [K^+^] is determined by

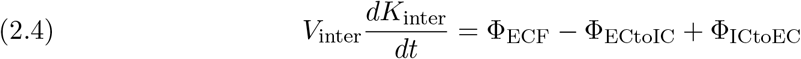

where *V*_inter_ is the interstitial fluid volume, Φ_ECtoIC_ and Φ_ICtoEC_ are the fluxes of K^+^ from the extracellular to the intracellular fluid and vice versa, respectively (see Eqs. 2.6 & 2.7). Lastly, *K*_IC_ is determined by the flux of K^+^ in and out of the cells

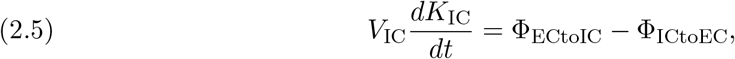

where *V*_IC_ denotes the intracellular fluid volume.

Flow of K^+^ into the intracellular fluid is driven by Na^+^-K^+^-ATPase uptake and is modeled using Michaelis-Menten kinetics so that

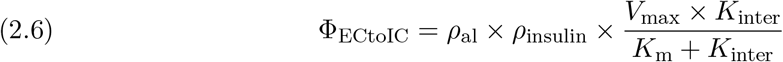

where *V*_max_ is the maximum rate, *K*_m_ denotes the half maximal activation level, *ρ*_al_ is the effect of ALD on Na^+^-K^+^-ATPase (see Section 2.1.3), and *ρ*_insulin_ is the effect of insulin (see Section 2.1.4). K^+^ returns to the extracellular compartment via diffusion through a permeable membrane:

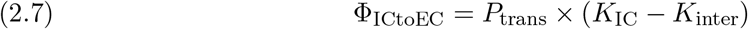

where *P*_trans_ denotes the transmembrane permeability.

#### 2.1.2. Renal K^+^ regulation

Filtered K^+^ load (Φ_filK_) is proportional to the GFR (Φ_GFR_) and plasma [K^+^] so that

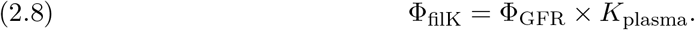

The filtrate then moves through the nephrons, where K^+^ is reabsorbed and secreted along the different segments. A schematic of the key segments for K^+^ renal transport is given in Fig. 2. At the end of the nephrons, what is remaining is excreted in urine. The model represents the kidney as a single representative nephron split into three segments: the “proximal segment (ps)” which includes the proximal tubule and the loop of Henle, the “distal segment (dt)” that includes the distal convoluted tubule and the connecting tubule, and the collecting duct (cd).

**Figure 2.**
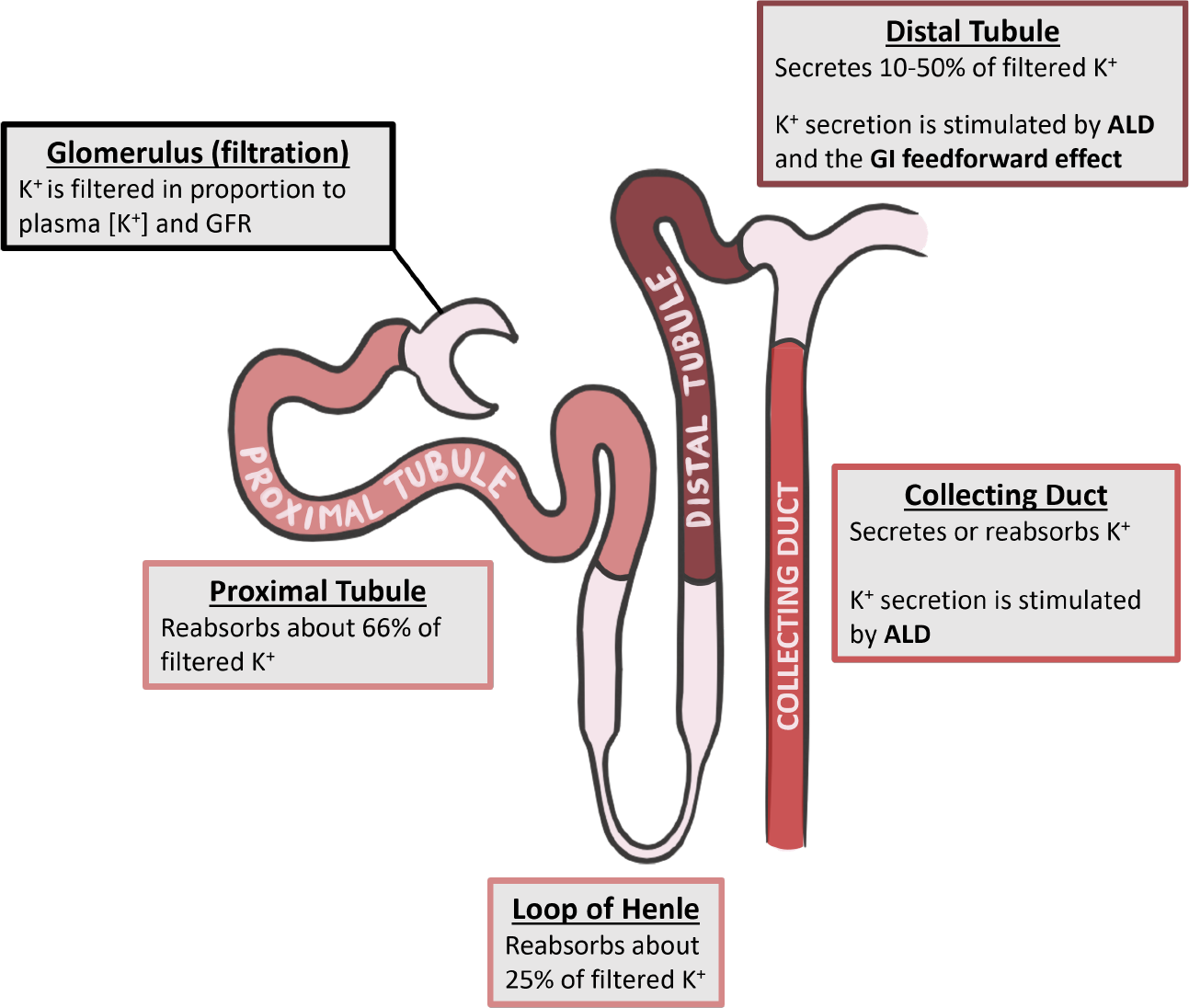
Filtration, reabsorption, and secretion of K^+^ in key nephron segments. Note that the proximal segment consists of the proximal tubule and the loop of Henle. Figure modified from Stadt et al. [19]. GFR: glomerular filtration rate; ALD: aldosterone; GI: gastrointestinal

Let *η*_ps*−*Kreab_ denote the fractional K^+^ reabsorption along the proximal segment so that net proximal segment K^+^ reabsorption is given by

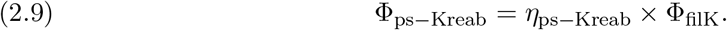

Distal tubule K^+^ secretion is modeled using a baseline distal tubule secretion value of 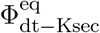 and is regulated by ALD and the gastrointestinal feedforward mechanism so that

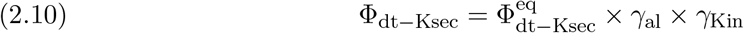

where *γ*_al_ denotes the regulatory effect of ALD (see Section 2.1.3) and *γ*_Kin_ represents the gastrointestinal feedforward effect (see Section 2.1.5). Similarly, collecting duct K^+^ secretion (Φ_cd*−*Ksec_) has a baseline value 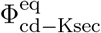 and is regulated by ALD so that:

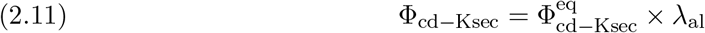

where *λ*_al_ denotes the regulatory effect of ALD (see Section 2.1.3). Collecting duct K^+^ reabsorption (Φ_cd*−*Kreab_) is modeled as a linear function depending on the filtrate entering the collecting duct based on transport from the previous segments:

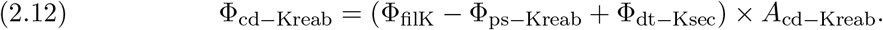

Finally, urine K^+^ excretion (Φ_uK_) is given by the filtration and the net transport along the various segments:

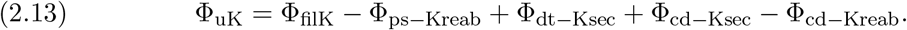

#### 2.1.3. Aldosterone effects

To model [ALD], denoted by *C*_al_, we use the approach developed by Maddah & Hallow [11]:

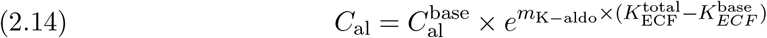

where *m*_K*−*aldo_ is a fitting parameter and 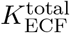 is the total extracellular fluid [K^+^] given by

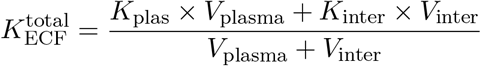

and 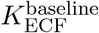 is the baseline extracellular [K^+^]. We capture the effect of [ALD] on Na^+^-K^+^-ATPase abundance by the scaling factor *ρ*_al_ (see Eq. 2.6), which is represented linearly as:

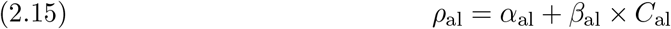

where *α*_al_ and *β*_al_ are parameters. The effect of [ALD] on distal tubule and collecting duct K^+^ secretion are represented by *γ*_al_ and *λ*_al_, respectively:

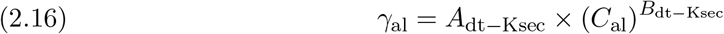

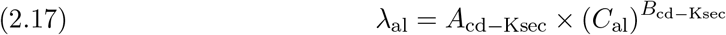

where *A*_dt*−*Ksec_, *B*_dt*−*Ksec_, *A*_cd*−*Ksec_ and *B*_cd*−*Ksec_ are fitting parameters.

#### 2.1.4. Insulin effects

The concentration of plasma insulin, denoted by *C*_insulin_ in the time after a meal is given by

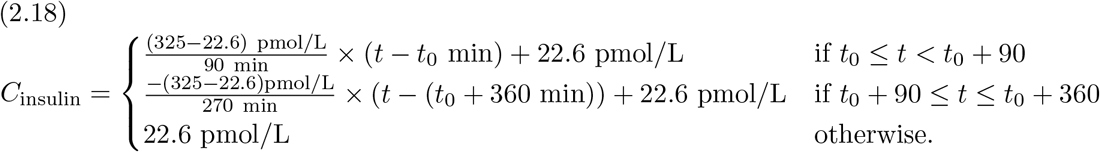

where *t*_0_ is the time at the beginning of the meal (see Ref. [19]). To model insulin stimulation of Na^+^*−*K^+^-ATPase, we let *ρ*_insulin_ (see Eq. 2.6) be given by

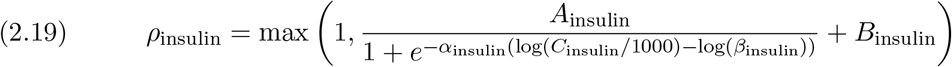

where *A*_insulin_ and *B*_insulin_ are fitting parameters.

#### 2.1.5. Gastrointestinal feedforward effect

The model represents gastrointestinal feed-forward effect via the term *γ*_Kin_, which alters distal tubule K^+^ secretion depending on *M*_Kgut_ (see Eq. 2.10) where

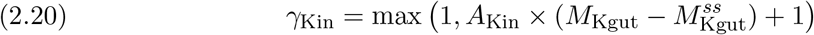

where *A*_Kin_ is a fitting parameter and 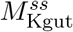 is the steady state value of M_Kgut_.

### 2.2. Morris method

The Morris method is a randomized one-at-a-time (OAT) sensitivity analysis method that has shown to be a computationally efficient efficient and reliable method for identifying and ranking important variables for complex systems [1, 14, 16]. A more comprehensive description of the Morris method is in Appendix B.

The Morris method involves computing elementary effects for each of the model factors by evaluating the model *r* times for different factor values. After computing the *r* elementary effects for each factor (denoted as the *i*-th factor), we compute the average value (*μ*_*i*_), average of the absolute values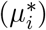, standard deviation (*σ*_*i*_), and Morris Index (*MI*_*i*_):

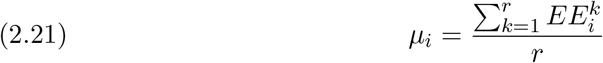

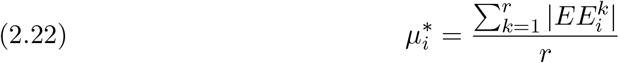

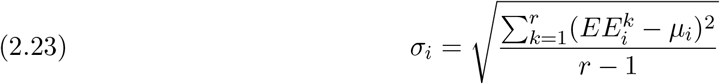

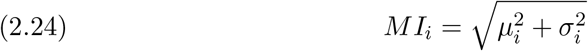

where 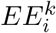 denotes the elementary effect of the *i*-th factor during the *k*-th model evaluation. We will refer to sensitivity analysis using the Morris method as a “Morris analysis” in short.

Results from a Morris analysis are often displayed in a Morris plot, where *σ*_*i*_ is plotted against 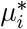 (or *μ*_*i*_) for each factor. The greater the value of 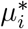 for factor *i*, the more the *i*-th factor affects the model output. The greater the *σ*_*i*_ value, the more the factor is nonlinear or involved with interactions with other factors; a low *σ*_*i*_, by contrast, indicates a linear, additive factor.

In our Morris analysis, the factors are the model parameters, with ranges given in Table 1. We chose *r* = 1000 for the number of model evaluations per parameter. An analysis is done separately for the steady state solutions and for the single meal simulations. For the steady state analysis, the we conduct the Morris analysis by sampling over the parameter space and computing the steady state solution for the baseline model for the sampled parameters with K^+^ intake (Φ_Kin_; Eq. 2.1) fixed at 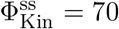. For the single meal simulations, we simulate the respective single meal simulation (see Section 2.3) with the sampled parameter set. The elementary effects are computed at simulation time points of every ten minutes because different parameters have differing effects at different time points in the simulation.

**Table 1.**
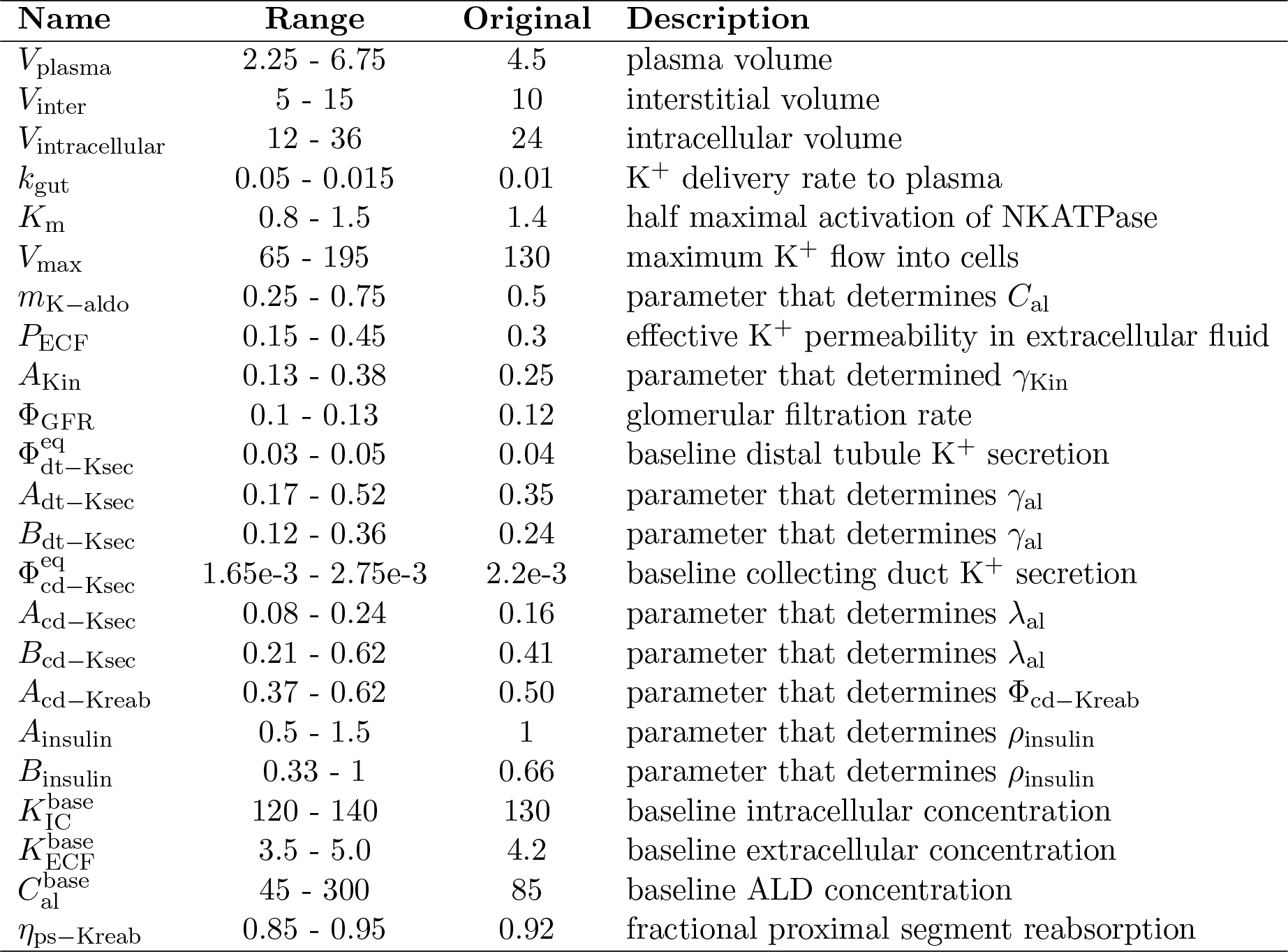
Parameter ranges and baseline values (used from Ref. [19]) used for the Morris analysis and baseline model simulations.

### 2.3. Single meal simulations

In our previous study (Ref. [19]), we investigated the physiological responses to meals of different contents by conducting *in silico* experiments of whole-body K^+^ regulation during a single meal of varying contents. To do this, we simulated three experiments based on the methods of the clinical trial presented in Preston et al. [15]. The experiments consisted of a meal deficient in K^+^ (“Meal only”), an oral ingestion of K^+^-Cl^*−*^ only (“KCl only”), and typical meal made up of a combination of the K^+^-deficient meal with K^+^ intake (“Meal + KCl”). Model simulations for the three meal types are shown in Fig. 3. The meal is given at time 0 after a fasting period of 6 hours from the steady state initial condition as done in Ref. [19]. In this study we refer to these simulations as the “single-meal simulations” and conduct a Morris analysis to determine the impacts of parameters on these results at time points of every 10 minutes in the simulation.

**Figure 3.**
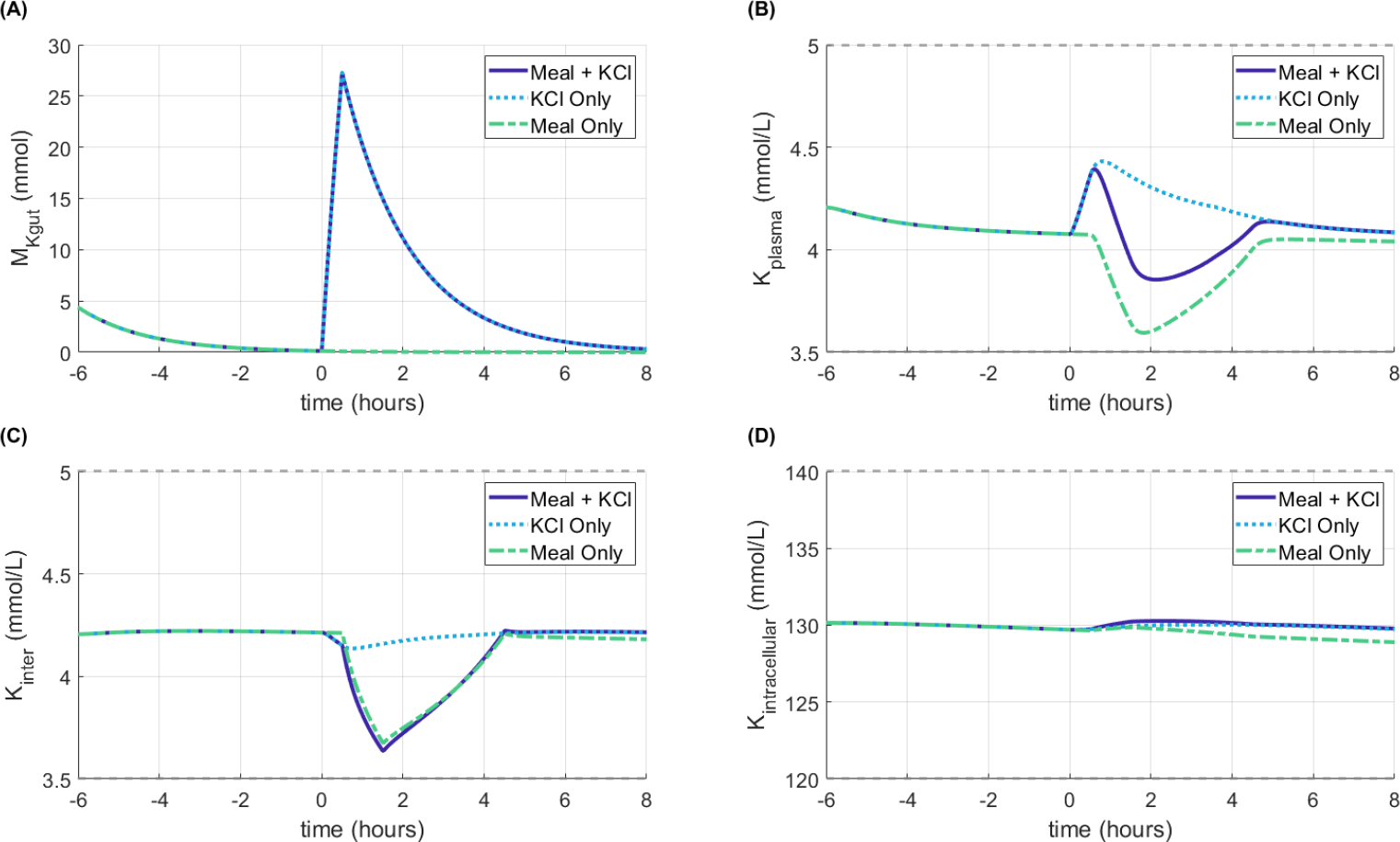
Model simulations using baseline parameters from Ref. [19] (listed in Table 1 as “Original”) for three meal simulation types: Meal + KCl, Meal only, and KCl only (Section 2.3). The meal starts at time = 0 after a 6 hour fasting period. Gray lines indicate normal range for (B) plasma, (C) interstitial, and (D) intracellular K^+^ concentration. The initial condition are the steady state solution. M_Kgut_: gastrointestinal K^+^ amount; K_plasma_: plasma K^+^ concentration, K_inter_: interstitial fluid K^+^ concentration; K_intracellular_: intracellular fluid K^+^ concentration

### 2.4. Software

The simulations and Morris analyses were conducted using `R`. We used the package `sensitivity` (https://cran.r-project.org/web/packages/sensitivity/index.html; Ref. [5]) to conduct the sensitivity analysis for the steady state and the package `ODEsensitivity` (https://cran.r-project.org/web/packages/ODEsensitivity/index.html; Ref. [20]) to conduct our sensitivity analysis for the single-meal simulations. The code used for this study is available at: https://github.com/Layton-Lab/Kreg_GSA.

## 3. Morris analysis results

### 3.1. Steady state analysis

To evaluate which parameters determine long-term K^+^ regulation, we conducted a Morris analysis on the model steady state. The Morris plots for parameter effects on model steady state plasma and intracellular [K^+^] are shown in Fig. 4. The Morris indices (Eq. 2.24) for plasma and intracellular [K^+^] are plotted in Fig. 5. Any parameters not included in the figures had inconsequential *μ*^*\**^, *σ*, and Morris index values for both plasma and intracellular [K^+^].

**Figure 4.**
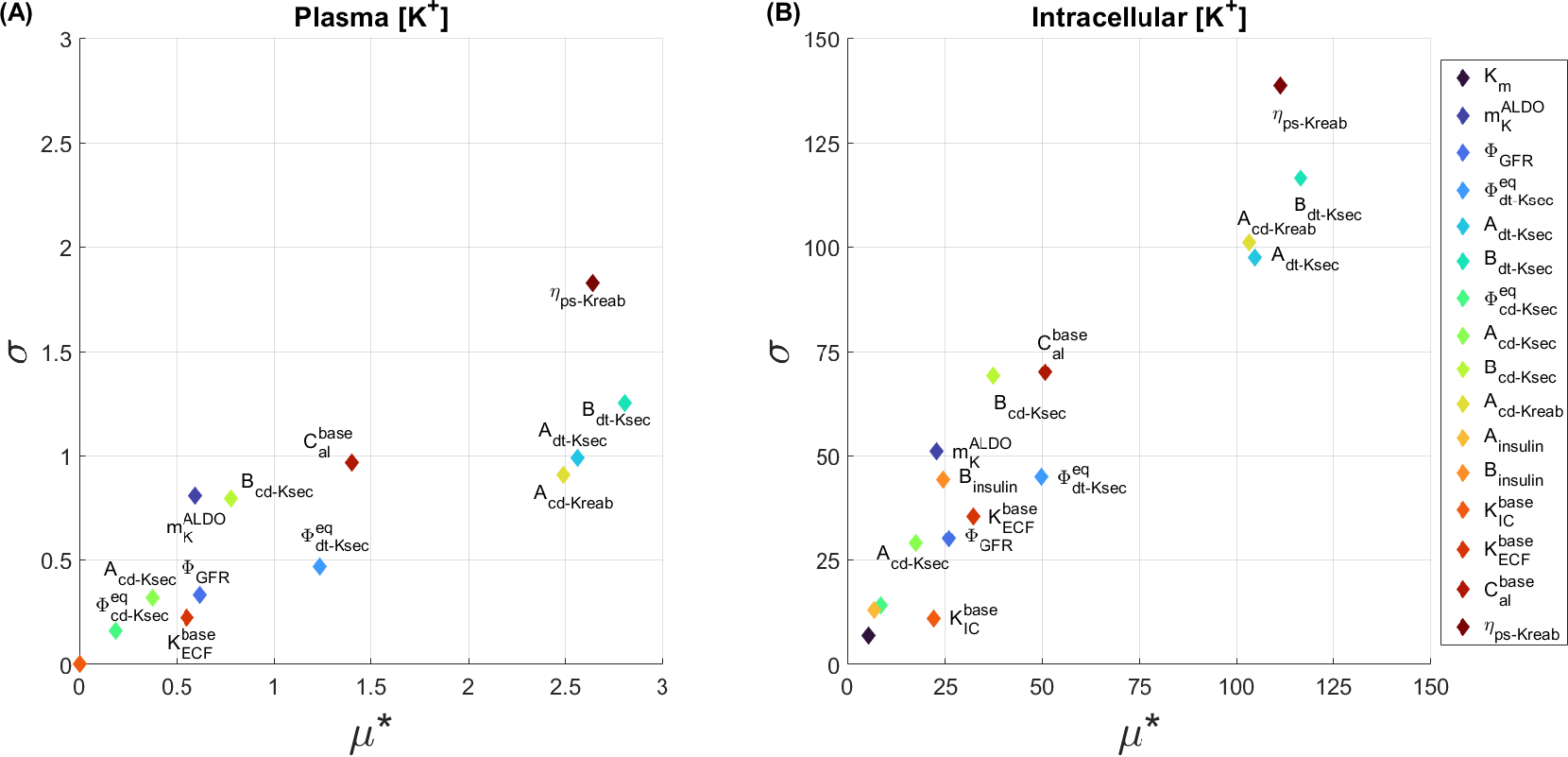
Morris plots for model steady state (A) plasma K^+^ concentration and (B) intracellular K^+^ concentration. Parameters not shown had μ^*^ and σ values at or near 0 for both plasma and intracellular K^+^ concentration.

**Figure 5.**
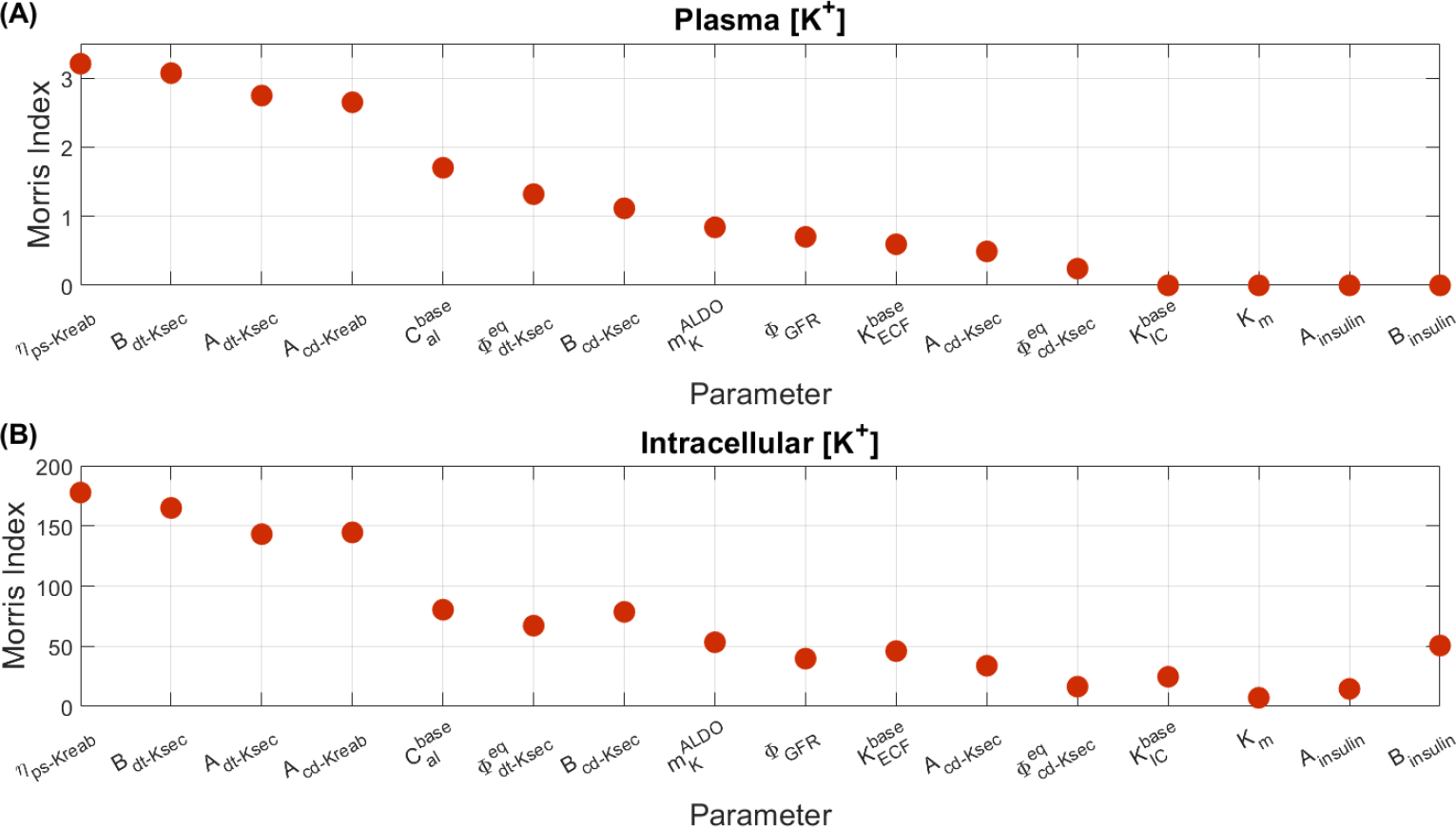
Morris indices (MI_i_; see Eq. 2.24) for model steady state (A) plasma K^+^ concentration and (B) intracellular K^+^ concentration. Parameters not shown had small Morris index values.

While the *μ*^*\**^, *σ*, and Morris index values are larger for the intracellular [K^+^] than plasma [K^+^], the eight largest *μ*^*\**^ values are the same parameters for both the plasma and intracellular [K^+^] (Fig. 4). Notably, the parameters with the top four Morris index values for both plasma and intracellular [K^+^] are *η*_ps*−*Kreab_, *B*_dt*−*Ksec_, *A*_dt*−*Ksec_, and *A*_cd*−*Kreab_, which are all parameters related to renal K^+^ handling (Fig. 5). Specifically, *η*_ps*−*Kreab_ represents fractional proximal segment K^+^ reabsorption (Eq. 2.9), *A*_dt*−*Ksec_ and *B*_dt*−*Ksec_ impact distal tubule K^+^ secretion (Eq. 2.16), and *A*_cd*−*Kreab_ impacts collecting duct K^+^ reabsorption (Eq. 2.12).

The parameter with the fifth highest Morris index is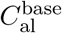, which represents baseline [ALD] (Fig. 5). Notably, the insulin parameters *A*_insulin_ and *B*_insulin_, which represent how insulin affects intracellular K^+^ intake (Eq. 2.19) do not have much impact on steady state plasma [K^+^] with a Morris index of about 0 (Fig. 5A). The insulin parameters do however have an impact on the intracellular [K^+^] (Fig. 5B), but are not the top parameters.

In summary, results of our Morris analysis indicate that the steady-state plasma and intracellular [K^+^] depend primarily on the renal handling parameters and baseline [ALD].

### 3.2. Single-meal simulations

The *μ*^*\**^ and *σ* values for parameter impact on plasma and intracellular [K^+^] during a single “Meal + KCl” simulation are shown in Fig. 6. During the fasting state (before time = 0), the *μ*^*\**^ and *σ* values do not change much for both plasma and intracellular [K^+^] from their initial value. When the meal is added, different parameters have a larger or smaller impact on model results, as determined by an increase or decrease in *μ*^*\**^ (Fig. 6A1-B1). For example, the *μ*^*\**^ values for *A*_insulin_ and *B*_insulin_, which determine insulin impact on Na^+^-K^+^-ATPase K^+^ uptake (Eq. 2.19), both increase. Additionally, *k*_gut_, which determines the delivery of K^+^ from the gastrointestinal system to the plasma (Eq. 2.1; Eq. 2.2) also increases from *μ*^*\**^*≈*0 to 0.5 for plasma [K^+^] (Fig. 6A1) and 0.64 for intracellular [K^+^] (Fig. 6B1) after the meal.

**Figure 6.**
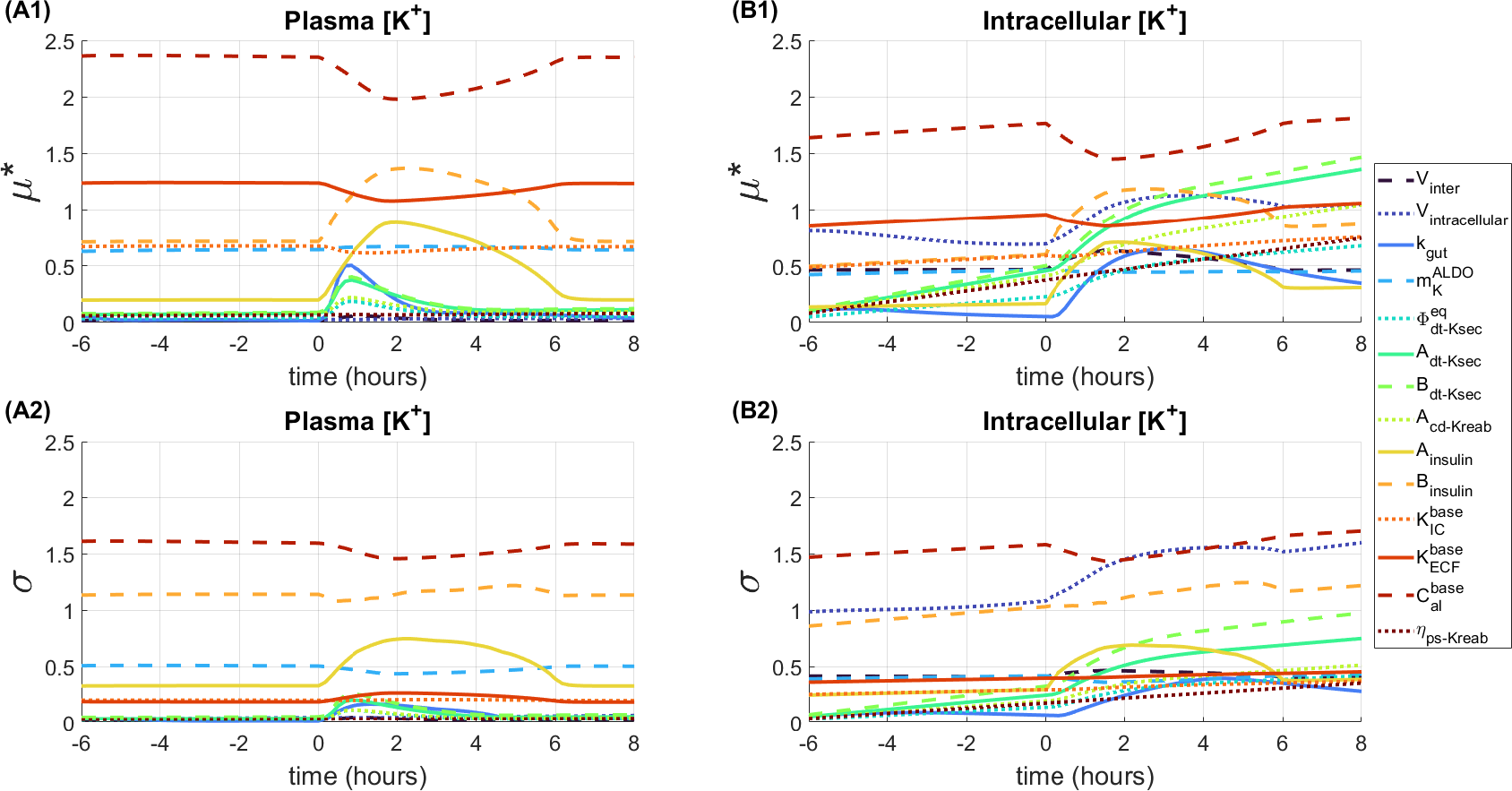
Morris (A1-B1) μ^*^ and (A2-B2) σ values for time points along the Meal + KCl simulation. Plasma [K^+^] effects are shown in panels (A1-A2) and intracellular [K^+^] effects are shown in (B1-B2). Elementary effects were compute for every 10 minutes of simulations. Only parameters that had significant μ^*^ and σ values through the simulations are shown.

Since the *μ* and *σ* values change over time in the simulation (as shown in Fig. 6), for clarity we computed the mean and standard deviation of the Morris indices over the full simulation for each parameter. These values are plotted with the minimum and maximum values in Fig. 7. For both plasma and intracellular [K^+^], 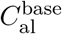 has the highest mean Morris index (Fig. 7). This is due to high *μ* and *σ* values for both plasma and intracellular [K^+^], showing that this parameter is both highly influential and interacts with other parameters.

**Figure 7.**
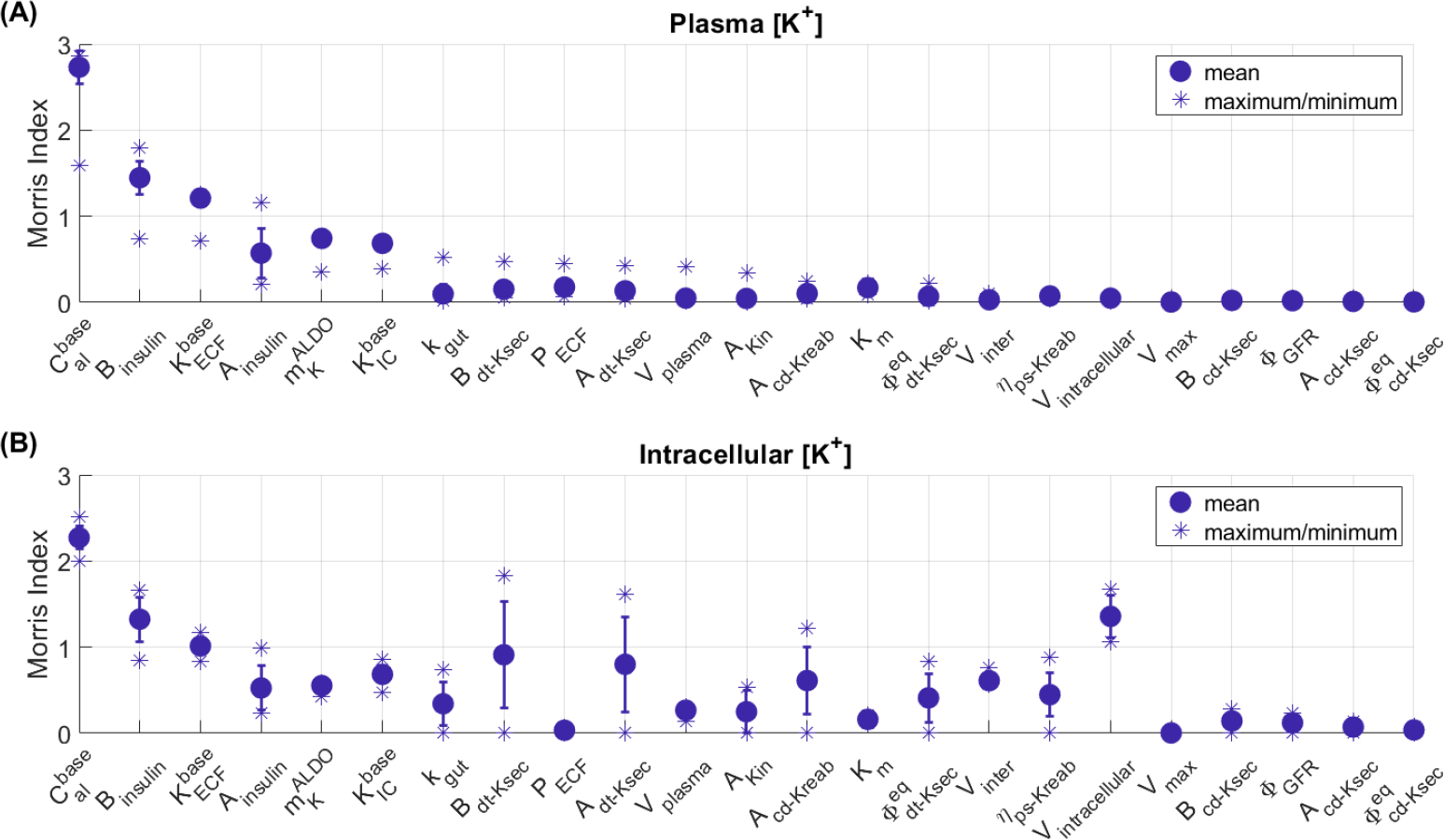
Mean, standard deviation (error bar), minimum, and maximum values of Morris indices along full Meal + KCl simulation for each parameter value for (A) plasma [K^+^] and (B) intracellular [K^+^].

Based on the maximum Morris index, the top five Morris indices for plasma [K^+^] are 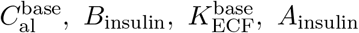, and 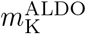. Note that these parameters are all a part of hormonal (i.e, ALD and insulin) signalling (Eq. 2.14; 2.19). Notably, for plasma [K^+^], *A*_insulin_ and *B*_insulin_ had Morris indices of nearly 0 in the steady state Morris analysis (Fig. 5A), but are in the highest Morris index values in the Meal + KCl simulation (Fig. 7A).

For intracellular [K^+^], the top five maximum Morris indices are 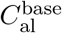, *B*_dt*−*Ksec_, *A*_dt*−*Ksec_, *V*_intracellular_, and *B*_insulin_ (Fig. 7B). Notably, the intracellular fluid volume (*V*_intracellular_) and ALD impact on distal tubule K^+^ secretion (shown by *A*_dt*−*Ksec_ and *B*_dt*−*Ksec_) were relatively insignificant for plasma [K^+^] (Fig. 7A) levels despite being in the parameters based on Morris indices for intracellular [K^+^] (Fig. 6B).

We conducted a Morris analysis for both the KCl only and Meal only single-meal simulations. The time series results are given in the Appendix A (Fig. 9; Fig. 10) and Morris indices are shown in Fig. 8. For KCl only, the results are about the same as the Meal + KCl results except that *A*_insulin_ and *B*_insulin_ have no impact since *ρ*_insulin_ = 1 (Eq. 2.19) in the KCl only simulation due to no insulin changes. Similarly, for Meal only, the results are similar to Meal + KCl except for a lower impact of *k*_gut_, *B*_dt*−*Ksec_, and *A*_dt*−*Ksec_ on intracellular [K^+^] (Fig. 8B) due to no addition of K^+^ into the system.

**Figure 8.**
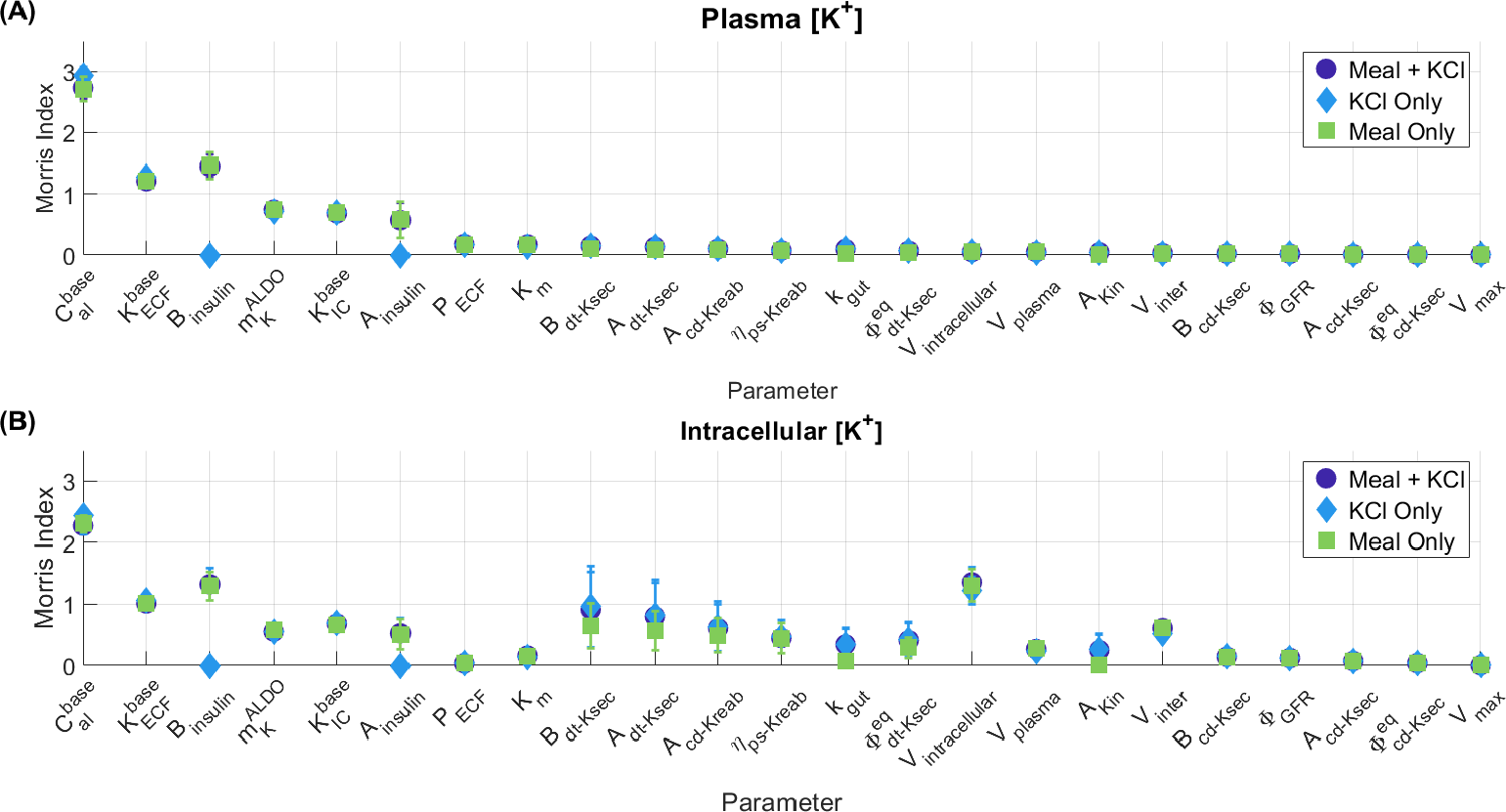
Mean and standard deviation of Morris indices along full Meal + KCl, KCl Only, and Meal Only simulation for each parameter value for (A) plasma [K^+^] and (B) intracellular [K^+^].

**Figure 9.**
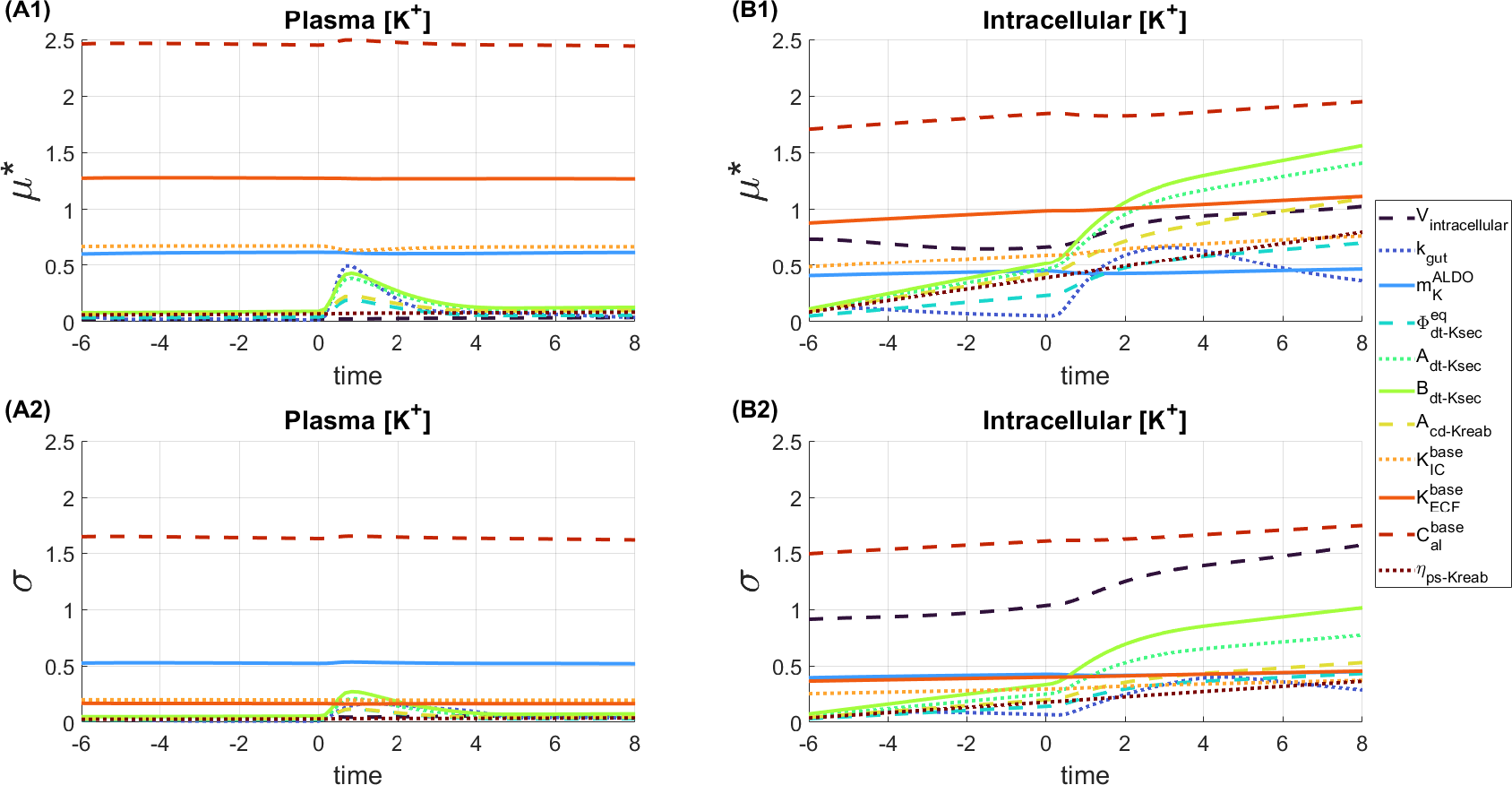
Morris (A1-B1) μ^*^ and (A2-B2) σ values for time points along the KCl Only simulation. Plasma [K^+^] effects are shown in panels (A1-A2) and intracellular [K^+^] effects are shown in (B1-B2). Elementary effects were compute for every 10 minutes of simulations. Only parameters that had significant μ^*^ and σ values through the simulations are shown.

**Figure 10.**
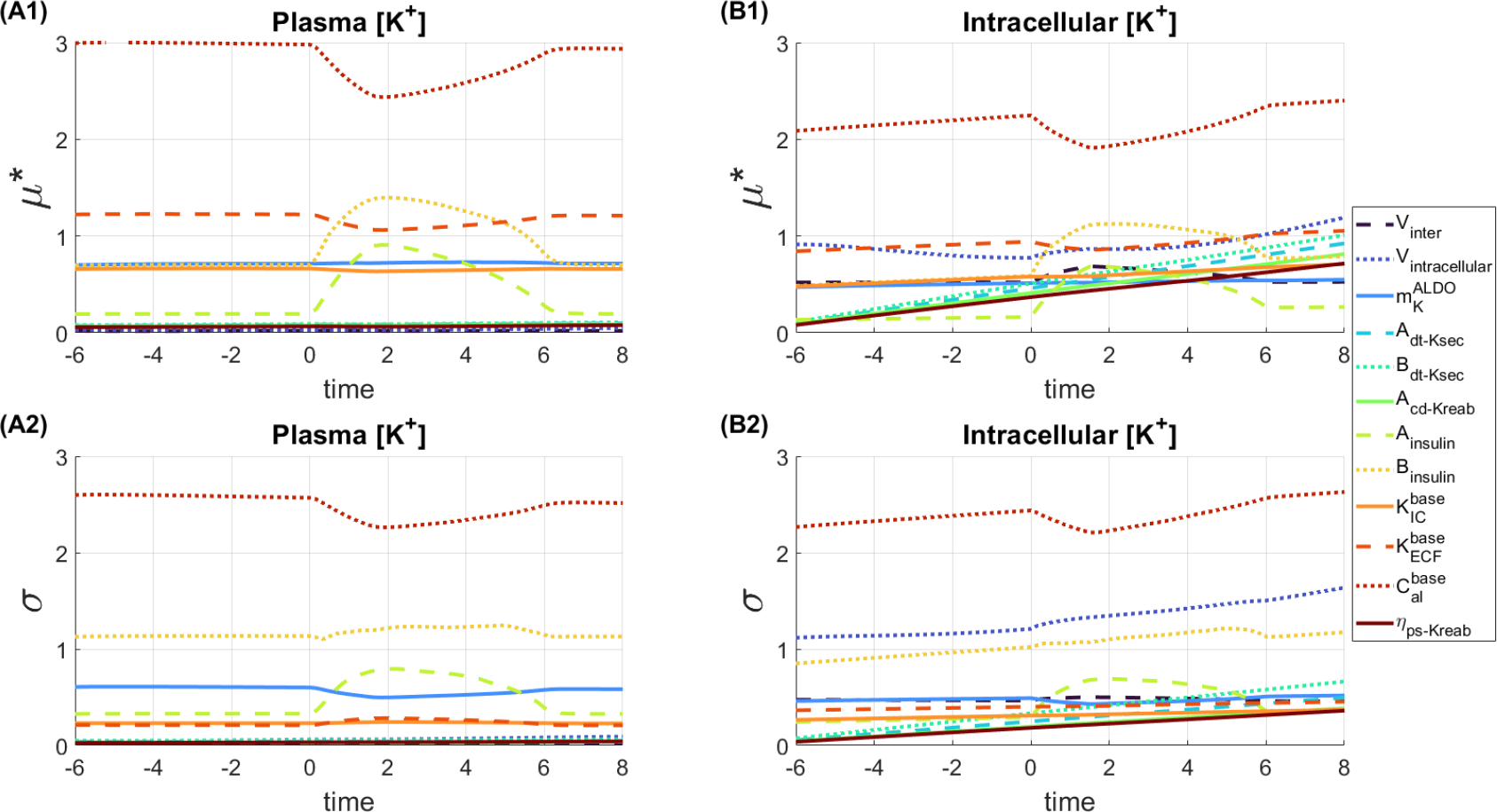
Morris (A1-B1) μ^*^ and (A2-B2) σ values for time points along the Meal Only simulation. Plasma [K^+^] effects are shown in panels (A1-A2) and intracellular [K^+^] effects are shown in (B1-B2). Elementary effects were compute for every 10 minutes of simulations. Only parameters that had significant μ^*^ and σ values through the simulations are shown.

## 4. Discussion

Sensitivity analysis can help identify sources of uncertainty in mathematical models. Models of complex systems, such as the K^+^ regulation model presented here, often involve a large number of parameters, many of which have unknown or uncertain values [17, 18]. There are two types of uncertainty in models of complex systems: epistemic uncertainty and aleatory uncertainty [17]. Epistemic uncertainty arises from incomplete knowledge of a system. This is often the case because parameters that ideally would be measured are often expensive or even impossible to measure in many systems. For example, measuring the exact amount that distal tubule K^+^ secretion increases for increased [ALD] would be an extremely invasive experiment. Aleatory uncertainty refers to the uncertainty that comes from the inherent heterogeneity in the system [17]. Indeed, making a model and choosing a single parameter set that represents the mean of a population would not represent each individual person. Rather, all people have differences in their body inter-individually as well as different states due to varying factors such as diet, exercise, age, and time of day that influence how an individual regulates K^+^. For this reason, studying the effects of varied parameters using global sensitivity analysis methods can help us further understand uncertainties that arise when modeling complex systems.

In this study, we conducted a sensitivity analysis of a mathematical model of whole-body K^+^ regulation (Ref. [19]) to understand how model parameter values may impact the model steady state and single-meal simulation predictions. Overall, our model analysis reveals that the K^+^ regulation model captures what is expected biologically: steady state plasma [K^+^] regulation is driven by renal function and plasma [K^+^] regulation after a shift in plasma [K^+^] (i.e., a single-meal) is driven by hormonal signalling. This analysis provides evidence that our mathematical model of whole-body K^+^ regulation (Ref. [19]) captures how the body regulates K^+^ both at homeostasis and during periods of K^+^ shifts.

### 4.1. Steady state results depend on renal handling of K^+^

In the Morris analysis of the model steady state, the top four parameters that the steady state plasma and intracellular [K^+^] are both most sensitive to are involved in renal K^+^ handling: *η*_ps*−*Kreab_, *B*_dt*−*Ksec_, *A*_dt*−*Ksec_, *A*_cd*−*Kreab_ (Fig. 4). Indeed, physiologically, long-term K^+^ handling is controlled by the kidneys, which excrete excess K^+^ through urine.

The *μ*^*\**^, *σ*, and Morris index values are larger for the intracellular than plasma [K^+^] in the steady state Morris analysis (Fig. 4; Fig. 5). This makes sense biologically because baseline plasma [K^+^] levels are around 3.5-5.0 mmol/L and baseline intracellular [K^+^] is about 120-140 mmol/L. Therefore intracellular [K^+^] levels have a wider possible range in value. Additionally, biologically, plasma [K^+^] levels are controlled more tightly, with intracellular [K^+^] fluctuating more to ensure a safe plasma [K^+^] [13]. Our model more tightly regulates plasma [K^+^] levels than intracellullar [K^+^] levels.

### 4.2. Aldosterone has a large impact on K^+^ regulation as well as interactions with other parameters

The parameter 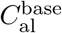, which represents baseline [ALD], was a top parameter for plasma and intracellular [K^+^] in both the steady state Morris analysis and the single-meal experiments. ALD is known to play a key role in K^+^ homeostasis. Notably, normal [ALD] levels have a wide range (about 45-300 ng/L; Ref. [3, 11]). We allowed 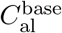 to vary by this known physiological range (Table 1), which is the largest range of all the parameters. This is likely a reason that this parameter had such large *μ*^*\**^ and *σ* values. However, we investigated the Morris analysis with a smaller range for this parameter and it still was a top parameter (results not shown).

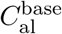 had a high variance in the elementary effects values, as given by large values for *σ*, especially in the single-meal experiments (Fig. 6). This indicates that this parameter has large interactions with other parameters. This makes sense biologically because someone who has typically low [ALD] level might respond differently to higher levels of [ALD] than someone with typically higher [ALD]. Indeed, our results show that, in isolation, [ALD] levels alter K^+^ regulation significantly.

### 4.3. Plasma K^+^ regulation after a single-meal is driven by hormonal signaling

The insulin parameters *A*_insulin_ and *B*_insulin_, which represent how insulin affects intracellular K^+^ uptake, do not have much impact on steady state plasma [K^+^] with a Morris index of about 0 (Fig. 5A). However, in the single-meal experiments, *B*_insulin_ and *A*_insulin_ became significant parameters for plasma [K^+^] with increasing *μ*^*\**^ and *σ* values after the meal, when insulin levels are increased (Fig. 6A1-A2). Based on the maximum Morris index, *B*_insulin_ and *A*_insulin_ were in the top 5 parameters for plasma [K^+^] in the single-meal experiments that included insulin (i.e., Meal + KCl and Meal only; Fig. 8A).

Additionally, for the single-meal simulations 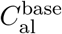 and 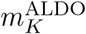, which are both parameters related to [ALD] (Eq. 2.14), are in the top 5 parameters for plasma [K^+^] (Fig. 8A). Therefore, our model captures that after a single-meal, which causes a shift in plasma [K^+^], is regulated primarily by hormonal signalling.

In conclusion, the Morris analysis results of our study capture the expected biological behaviour of K^+^ regulation under homeostasis conditions (i.e., steady state) and after a single meal of varying types. Results of the analysis can be used to inform future modeling development decisions, e.g., which parameters may best capture disease state models or therapeutic treatments.

## Appendix A. Additional Figures

## Appendix B. Morris method description

Let *f* be a mathematical model where the output is given by *f* (*X*_1_, *X*_2_, …, *X*_*n*_) and *X*_1_, *X*_2_, …, *X*_*n*_ are the model inputs, which will be referred to as factors. When computing a sensitivity analysis using the Morris method, we start by determining a defined range for all the possible values for the factors. The factors are then rescaled to be uniformly distributed on the unit interval and an initial base value is selected at random from this distribution. Subsequently, one random factor, *X*_*i*_, is incremented by a step size Δ, typically chosen to be *n/*(2(*n −* 1)), where *n* is the number of factors. The elementary effect for the *i*-th factor, denoted by *EE*_*i*_ is computed by:

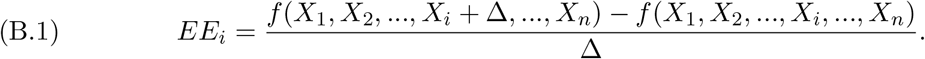

From this next value another random factor is incremented and the subsequent elementary effect is computed, until an elementary effect for each factor has been determined. This process is repeated *r* times by sampling at different points in the factor space. As a result, there will be a total of *r* elementary effects per factor at the end of the computations.

After computing the elementary effects for each factor, we can find the average value (*μ*_*i*_), average of the absolute values (*μ*^*\**^), standard deviation (*σ*), and Morris Index (*MI*_*i*_):

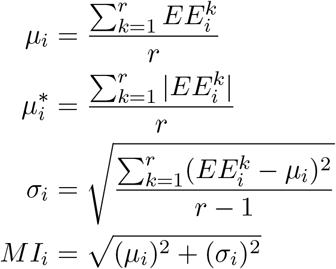

where 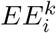 denotes the elementary effect of the *i*-th factor during the *k*-th model evaluation. The metrics can be interpreted in the following ways:

- *μ*^*\**^ gives a factor ranking: a greater 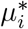 value indicates that the *i*-th factor affects the model output
- *σ* indicates that this factor interacts with other factors or is nonlinear
- *MI*: The Morris index is another way to factor rank by giving a single metric that incorporates both the mean (*μ*) and standard deviation (*σ*)

In an ordinary differential equation (ODE) model scenario, the factors are typically the model parameters. The model output measured can taken as the steady state solution, the values of the state variables at given time points in a simulation, or another measure that can be computed from a given model simulation.

## Notes

**Funding:** This work was supported by the Canada 150 Research Chair program and by the Natural Sciences and Engineering Council of Canada, via a Discovery award (RGPIN-2019-03916).

https://github.com/Layton-Lab/Kreg_GSA

### Competing Interest Statement

The authors have declared no competing interest.

